# *Aspergillus fumigatus* hypoxia adaptation is critical for the establishment of fungal keratitis

**DOI:** 10.1101/2023.10.01.560368

**Authors:** Jorge D. Lightfoot, Emily M. Adams, Manali M. Kamath, Becca L. Wells, Kevin K. Fuller

## Abstract

Purpose: The poor visual outcomes associated with fungal keratitis (FK) underscore a need to identify fungal pathways that can serve as novel antifungal targets. In this report, we investigated whether hypoxia develops in the FK cornea and, by extension, if fungal hypoxia adaptation is essential for virulence in this setting. Methods: C57BL/6j mice were inoculated with *Aspergillus fumigatus* and *Fusarium solani* var *petroliphilum* via topical overlay or instrastromal injection. At various time points post-inoculation (p.i.), animals we were injected with pimonidazole for the detection of tissue hypoxia through immunofluorescence imaging. The *A. fumigatus srbA* gene was deleted through Cas9-mediated homologous recombination and its virulence was assessed in the topical infection model using slit-lamp microscopy and optical coherence tomography (OCT). Results: Topical inoculation with *A. fumigatus* resulted in diffuse pimonidazole staining across the epithelial and endothelial layers within 6 h. Stromal hypoxia was evident by 48 h p.i., which corresponded to leukocytic infiltration. Instrastromal inoculation with either *A. fumigatus* or *F. solani* similarly led to diffuse staining patterns across all corneal cell layers. The *A. fumigatus srbA* deletion mutant was unable to grow at oxygen levels below 3% *in vitro,* and corneas inoculated with the mutant failed to develop signs of corneal opacification, inflammation or fungal burden. Conclusions: These results suggest that fungal antigen rapidly drives the development of corneal hypoxia, thus rendering fungal SrbA or related pathways essential for the establishment of infection. Such pathways may therefore serve as targets for novel antifungal intervention.

## Introduction

*Aspergillus fumigatus* is a ubiquitous mold and an important agent of human infection. It is the predominant fungal pathogen of the lung, for example, where it can cause an allergic bronchopneumonia in atopic individuals or an invasive infection in those with severely compromised immune systems^1,2^. The latter disease, termed invasive pulmonary aspergillosis (IPA), carries a mortality rate of 40-90% depending on the extent of immune suppression or fungal dissemination to distal organs^3,4^. *A. fumigatus* is also an important cause of fungal keratitis (FK), a sight threatening infection of the cornea that can affect otherwise healthy individuals that are inoculated through ocular trauma or contact lens wear^5,6^. FK results in corneal perforation or the need for corneal transplantation in 40-60% of cases and long-term vision impairment in up to 70% of patients^7^. The poor patient outcomes in either IPA or FK is due, in part, to the inadequacy of currently available antifungals^8–10^. In principle, fungal proteins or signaling pathways that promote adaptation and growth within the host tissue could serve as novel drug targets, but the degree to which pathways differ as a function of host site is poorly understood. Accordingly, we seek to identify fungal pathways that support virulence in both the lung and corneal environments to facilitate the development of novel antifungals with dual use.

As the lung is perfused by ambient air (∼20% O_2_), it is perhaps unexpected that hypoxia develops during IPA. Nevertheless, Grahl *et al*. demonstrated that the chemical pimonidazole (Hypoxyprobe), which binds to proteins within hypoxic cells, accumulates within inflammatory foci of *A. fumigatus*-infected murine lungs several days after inoculation^11–13^. These low oxygen microenvironments (<1% O_2_) likely arise due to several factors, including a physical reduction of oxygen diffusion through the inflamed and necrotic tissue, as well as a rapid metabolic consumption of molecular oxygen by the inflammatory cells themselves^14,15^. Ethanol, which is not produced by host cells, is furthermore detectable in infected lungs at time points corresponding to the positive pimonidazole staining, indicating a shift in fungal metabolism towards fermentation^11^. Taken together, the cell compositional, structural, and metabolic changes that occur within the lung over the course of *A. fumigatus* infection cumulatively result in an oxygen depletion to which the fungus must adapt^11,16,17^.

A critical component of the *A. fumigatus* hypoxic response is the regulation of membrane sterol metabolism^18–20^. Like most fungi, *A. fumigatus* relies on the *de novo* synthesis of ergosterol to maintain membrane homeostasis and growth. The synthesis of each molecule of ergosterol is energetically costly, however, requiring a large amount of ATP equivalents and 12 molecules of oxygen^21,22^. This is significant in two related ways, the first being that as molecular oxygen becomes limiting, so too will the concentration of membrane ergosterol. Second, fungi lack an apparent ortholog of the mammalian Hif-1 alpha and cannot sense environmental oxygen directly^22^. Thus, the depletion of membrane sterols serves as proxy for hypoxia and triggers the adaptive response, which in *A. fumigatus* is chiefly driven by Zn-finger transcription factor SrbA (for <u>s</u>terol-<u>r</u>egulatory <u>e</u>lement-<u>b</u>inding <u>p</u>rotein)^20^. Upon sterol depletion, SrbA is cleaved by regulatory proteins, translocates from the ER membrane to the nucleus and, as the name suggests, drives the expression of several genes within the ergosterol biosynthetic pathway^19,23,24^. Whereas wild-type *A. fumigatus* grows almost uninhibited at oxygen concentrations as low as 0.2%, the *srbA* deletion mutant (Δ*srbA*) is growth ablated at oxygen concentrations at or below 3%^25^. The Δ*srbA* mutant is furthermore avirulent in multiple models of IPA, thus underscoring the importance of fungal hypoxia adaptation within the lung^20^.

It is currently unknown whether hypoxia develops in the fungal-infected cornea. On one hand, the primary site of fungal growth during FK, the central stroma, is separated from ambient air by a thin epithelium and a tear film that altogether measure just 550 µM in thickness in humans^26^. Thus, it may be expected that the cornea is unique among other host tissues in that it, and the infecting fungus, remain well-perfused by atmospheric oxygen during infection. On the other hand, the cornea lacks a vascular supply of oxygen and massive leukocytic infiltration and tissue edema are hallmarks of FK pathology^27^. These factors may therefore lead to oxygen depletion in the central cornea due to an increased diffusion barrier of atmospheric O_2_ (tissue thickness) and its conversion to ROS by host inflammatory cells, altogether mirroring the pathobiology of pulmonary aspergillosis^28–31^. In this report, we demonstrate that inoculation of murine corneas with *A. fumigatus* does indeed lead to the development of hypoxia, not just in the central cornea, but across all tissue layers including the epithelium. Interestingly, and unlike the lung, this hypoxia precedes the infiltration of peripheral leukocytes, suggesting that fungal antigen can influence oxygen metabolism in corneal resident cells. Importantly, we demonstrate that an *A. fumigatus srbA* deletant is unable to establish infection in our murine model, suggesting that the pathway could serve as a novel target for the treatment of FK.

## Results

### Corneal hypoxia develops in two distinct murine models of fungal keratitis

To determine if *A. fumigatus* infection promotes the development of corneal hypoxia, we first employed a topical inoculation model of FK previously described^32–34^. Briefly, 6-8-week-old C57BL/6j mice were immunosuppressed with methylprednisolone 24 h prior to infection. On the day of inoculation, an approximately 1 mm area of the central corneal epithelium was debrided with an Algerbrush and topically overlaid with *A. fumigatus* conidia that were pre-germinated (swollen, but not polarized) in nutrient rich broth. At 6, 12, 24, and 48 h post-inoculation (p.i.), animals were administered pimonidazole by intraperitoneal injection and euthanized 90 minutes later. Ocular sections were either stained with Periodic acid-Schiff and hematoxylin (PASH) for histological analysis, or with anti-pimonidazole antibodies for the detection of hypoxic microenvironments via immunofluorescence (IF). Whereas sham-inoculated and untouched (control) corneas were negative for pimonidazole staining at all tested time points, those inoculated with *A. fumigatus* displayed positive staining in the epithelium, endothelium and even discrete layers of the stromal collagen beginning at the earliest tested time point (6 h p.i.). The central epithelial ulcer generated for the inoculation typically reforms within 24 h in un-inoculated corneas, but does not reform in infected corneas; consequently, the sections shown in Figure 1 are peripheral to the original ulcer (i.e., closer to the limbus). The distribution and intensity of the pimonidazole staining was markedly enhanced at 48 h p.i., which corresponded to both the presence of leukocytic infiltration and tissue damage upon evaluation of PASH stained sections as well as the onset of corneal opacification upon slit-lamp examination (**Figure 1**). Taken together, the topical inoculation of corneas with *A. fumigatus* results in the rapid development of hypoxia that precedes, but is exacerbated by, the influx of inflammatory cells into the tissue.

**Figure 1.**
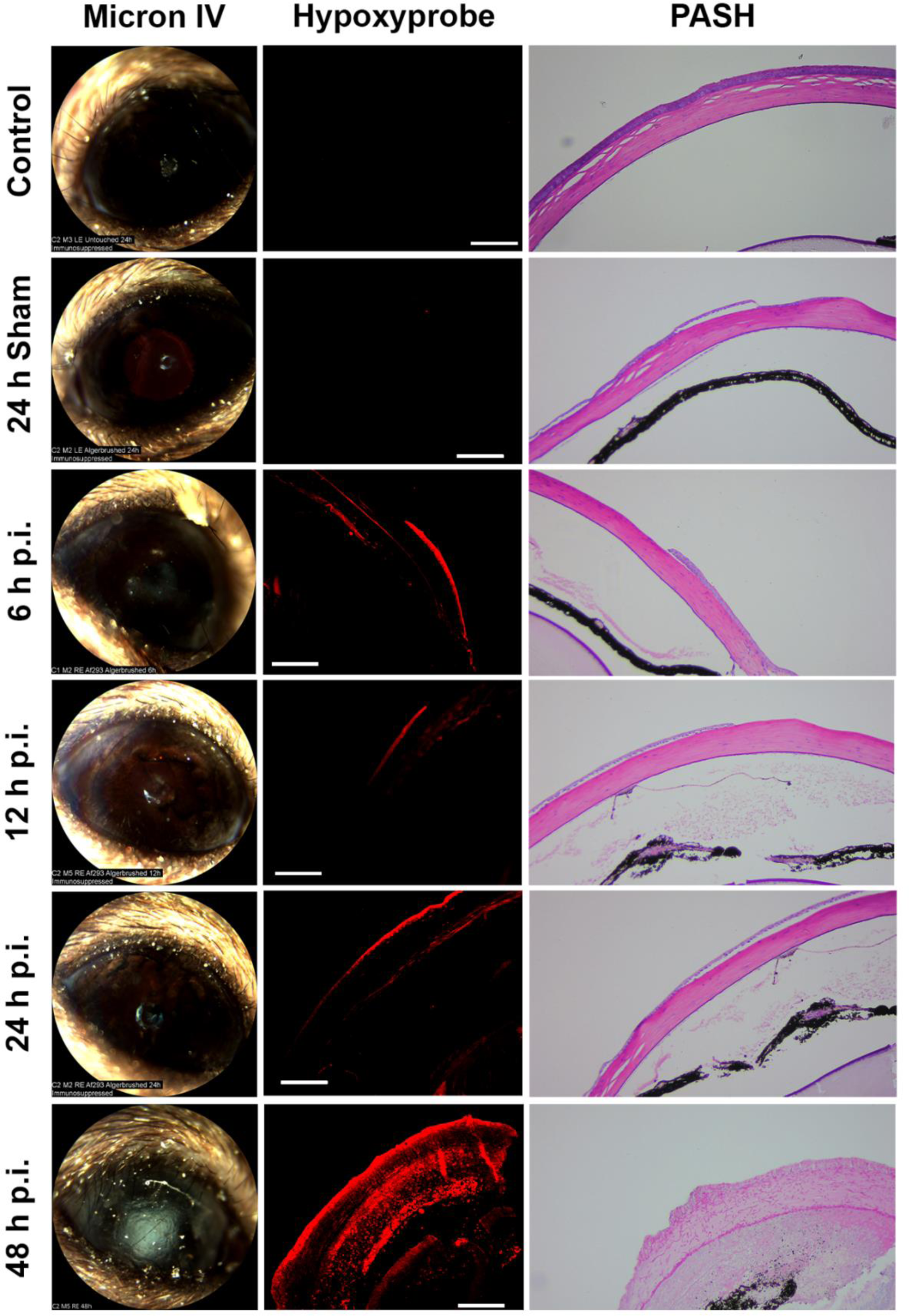
A topical model of infection causes the development of hypoxia in 6 hours p.i. as observed by staining immunohistochemically with Hypoxyprobe. In a topical model of infection, Hypoxyprobe was injected intraperitoneally 90 minutes before the endpoints at 6, 12, 24, and 48 hours p.i.. Control and Sham infected eyes were taken at 24 h p.i.. Eyes were processed histologically and an anti-Hypoxyprobe antibody was used to detect Hypoxyprobe bound to proteins (red) in one section, the following section was Periodic Acid Schiff and Hematoxylin (PASH) stained for the presence of *A. fumigatus* (fuchsia). Isotype controls and mice not injected with Hypoxyprobe show that staining was specific. (shown in SI1). Scale bars are 100 µm.

We reasoned that the inoculation onto the corneal surface in the above study could lead to hypoxia across the epithelium by at least two, but possibly additive ways: first, the direct interaction of concentrated fungal antigen with epithelial cells could lead to altered metabolism and oxygen consumption by those cells; second, the fungus itself may consume and locally deplete the oxygen availability to the tissue layer. We hypothesized that in either case, such patterns of epithelial hypoxia would be absent if direct fungal-epithelial cell interactions were largely bypassed. To explore this idea, we next evaluated patterns of pimonidazole staining following an intrastromal injection method, whereby *A. fumigatus* conidia were delivered directly into the underlying stroma through a small needle puncture. In parallel, we also inoculated animals with another predominant FK pathogen, *Fusarium solani* var *petroliphilum,* to determine if hypoxia development was pathogen-specific. In contrast to the technical ease and high infection rate associated with topical inoculation, we experienced a high failure rate (∼50%) for the intrastromal approach that that was not obvious until animals did or did not develop signs of disease at around 24 h p.i. As such, early (sub-clinical) time points could not be confidently analyzed as they were with the topical infection model. Nevertheless, animals that did develop signs of corneal opacification at 24 h p.i. displayed not only a positive pimonidazole signal in the intrastromal space, as expected, but also a strong signal across the entire corneal epithelial and endothelial layers similar to what was observed following topical inoculation (**Figure 2**). Unstained infected tissue sections or those stained with isotype control antibodies did not display a positive fluorescent signal (**Figure S1**). Taken together, hypoxia development across the corneal cell layers was dependent neither on the mode of fungal inoculation nor the pathogen evaluated in this study.

**Figure 2.**
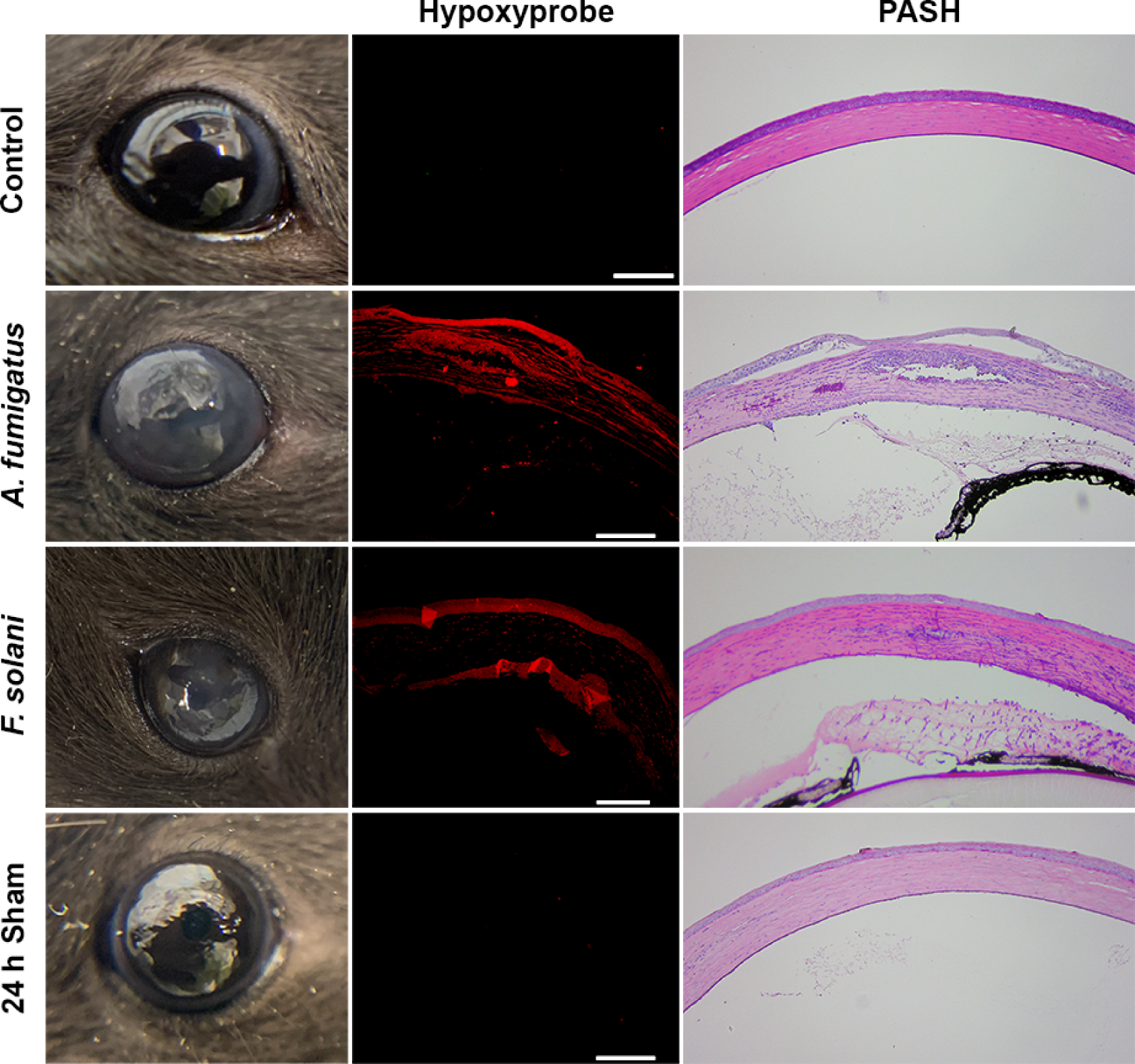
Hypoxia develops during intrastromal infection at 24 hours p.i.: 5×10^4^ Af293 conidia are injected into the corneal stroma of mice immunosuppressed with methylprednisolone. The infected corneas stained positively for Hypoxyprobe diffusely throughout the stroma and in the epithelium. In the following section stained with PASH, the presence of fungal burden (fuchsia) is observed in punctate lesions alongside granulocytes (purple) in the stroma. The sham infected eye showed no Hypoxyprobe signal or fungal burden. Scale bars are 100 µm.

### The SrbA transcription factor is required for growth of *A. fumigatus* under hypoxia, but not on proteinaceous substrates

We next wanted to determine if the hypoxia that develops during FK has an influence on the pathobiology of the fungus. Towards this end, the gene encoding the *srbA* transcription factor was replaced with the phleomycin resistance cassette in our mCherry expressing *A. fumigatus* strain (strain Af293 *PgpdA-mCherry-hph*) using Cas9-mediated homologous recombination. Subsequent generation of the complemented strain (*ΔsrbA* C’) was achieved through the targeted integration of the wild-type *srbA* allele into the *atf4* locus of the *ΔsrbA* mutant (**Figure S2**)^35^. The isogenic strains were indistinguishable on glucose minimal medium (GMM), containing glucose and ammonium, under atmospheric conditions of 21% O_2_ (normoxia). By contrast, *ΔsrbA* was growth ablated on GMM at O_2_ concentrations below 3%, thus confirming the essential role for SrbA in *A. fumigatus* hypoxia adaptation previously described (**Figure 3A & 3B**).

**Figure 3.**
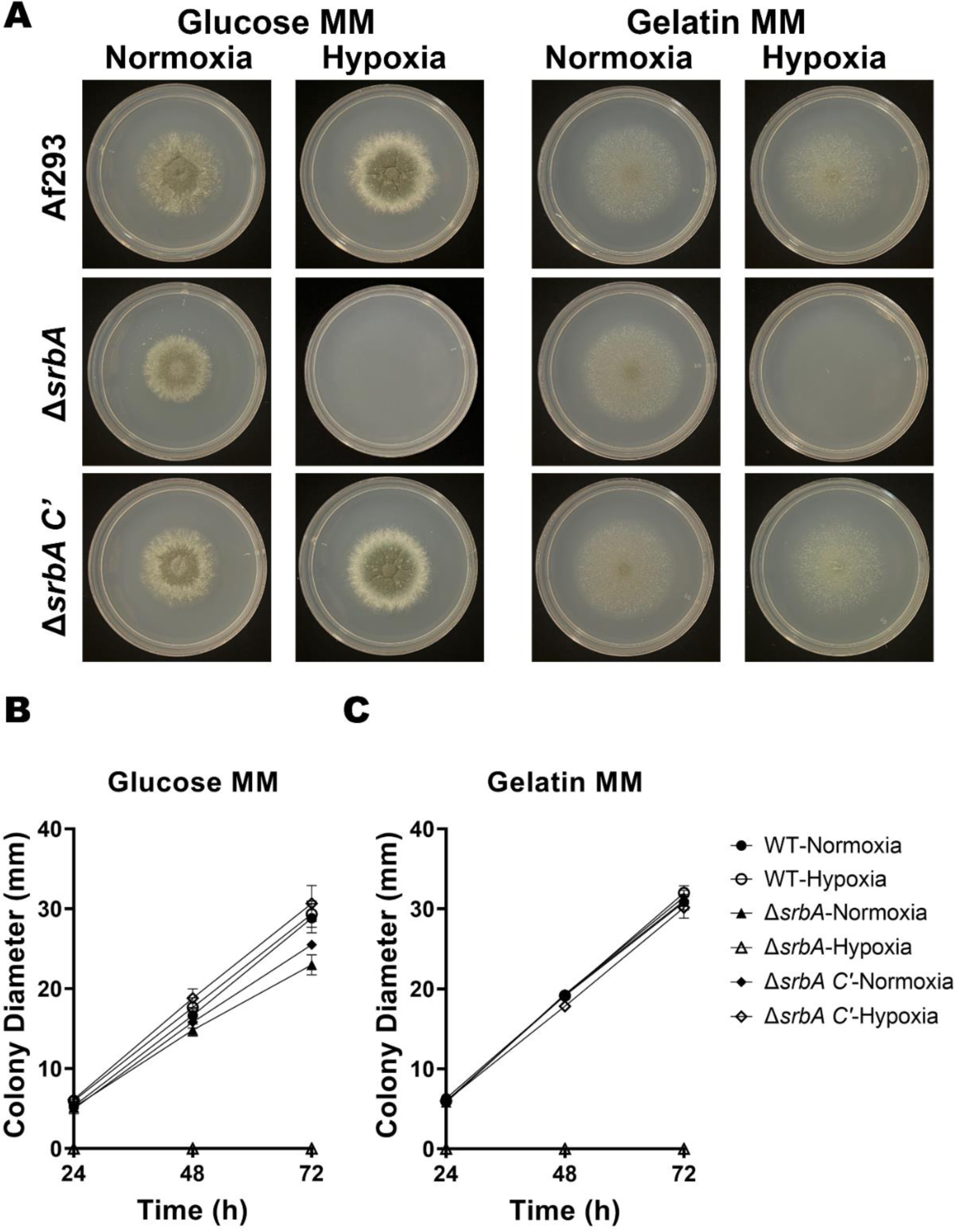
SrbA regulates the hypoxic signaling response in *A. fumigatus.* (A). The SrbA-null mutant was unable to grow at all in hypoxia (<1% O_2_), however it grew comparably to wild type in normoxia at 72 h post-inoculation on a range of media, including Glucose Minimal Media (B) and Gelatin Minimal Media (C). Diameters were measured every 24 h post-inoculation for 72 h. Significance measured by Kruskal-Wallis test.

Given that the cornea is rich in alternative macronutrients, primarily collagen and glycosaminoglycans that comprise the stromal extracellular matrix, we further tested the growth of the strains on media containing gelatin (collagen hydrolysate) as the sole carbon and nitrogen source. As shown in Figure 3, all strains were indistinguishable with respect to colony morphology and linear growth rate in normoxia. Similar to glucose, however, *ΔsrbA* was growth ablated in hypoxia on this medium (**Figure 3A & 3B**). We reasoned, therefore, that any observed virulence defect of *ΔsrbA* could be attributed to the influence of oxygen depletion in the tissue, rather than some previously uncharacterized role for SrbA in the metabolism of cornea-relevant macronutrients.

### *A. fumigatus* SrbA is essential for the establishment of fungal keratitis

We next tested the relative capacity of the strains to infect and drive disease in the above-described topical model of FK. In contrast to the WT and *srbA C’* groups, animals inoculated with Δ*srbA* failed to demonstrate any signs of disease upon external (slit-lamp) evaluation (**Figure 4A & 4B)**. Alterations in corneal structure were evaluated more thoroughly with optical coherence tomography (OCT), which provides a cross-sectional image of the anterior segment in live animals that can then be used for a quantitative assessment of corneal thickness. As expected, OCT revealed thickened corneal tissue and refraction in WT and *srbA C’*-infected corneas, which was indicative of the corneal edema and inflammation that are characteristic of FK. By contrast, and consistent with the external images, corneas inoculated with *ΔsrbA* were indistinguishable from sham-inoculated controls (**Figure 4C**). Tissue sections taken at 72 h p.i. were consistent with the OCT findings and demonstrated that WT and *srbA C’*-infected corneas were marked by ulcerated and structurally abnormal corneas, with massive immune cell infiltration in both the cornea and/or anterior chamber. Fungal hyphae were also observed through the depth of the cornea in these two groups on histology, and colony forming unit (CFU) assessment from homogenized corneas indicated comparable fungal loads. Histology of *ΔsrbA*-infected corneas, on the other hand, displayed normal epithelial and stromal architecture, minimal leukocytic infiltration, and had no visible fungal growth, the latter which was confirmed upon CFU analysis (**Figure 4D**). These results, which support a critical role of *A. fumigatus* SrbA in the establishment of corneal infection, were replicated in an independent experiment using both male and female animals (**Figure S3**).

**Figure 4.**
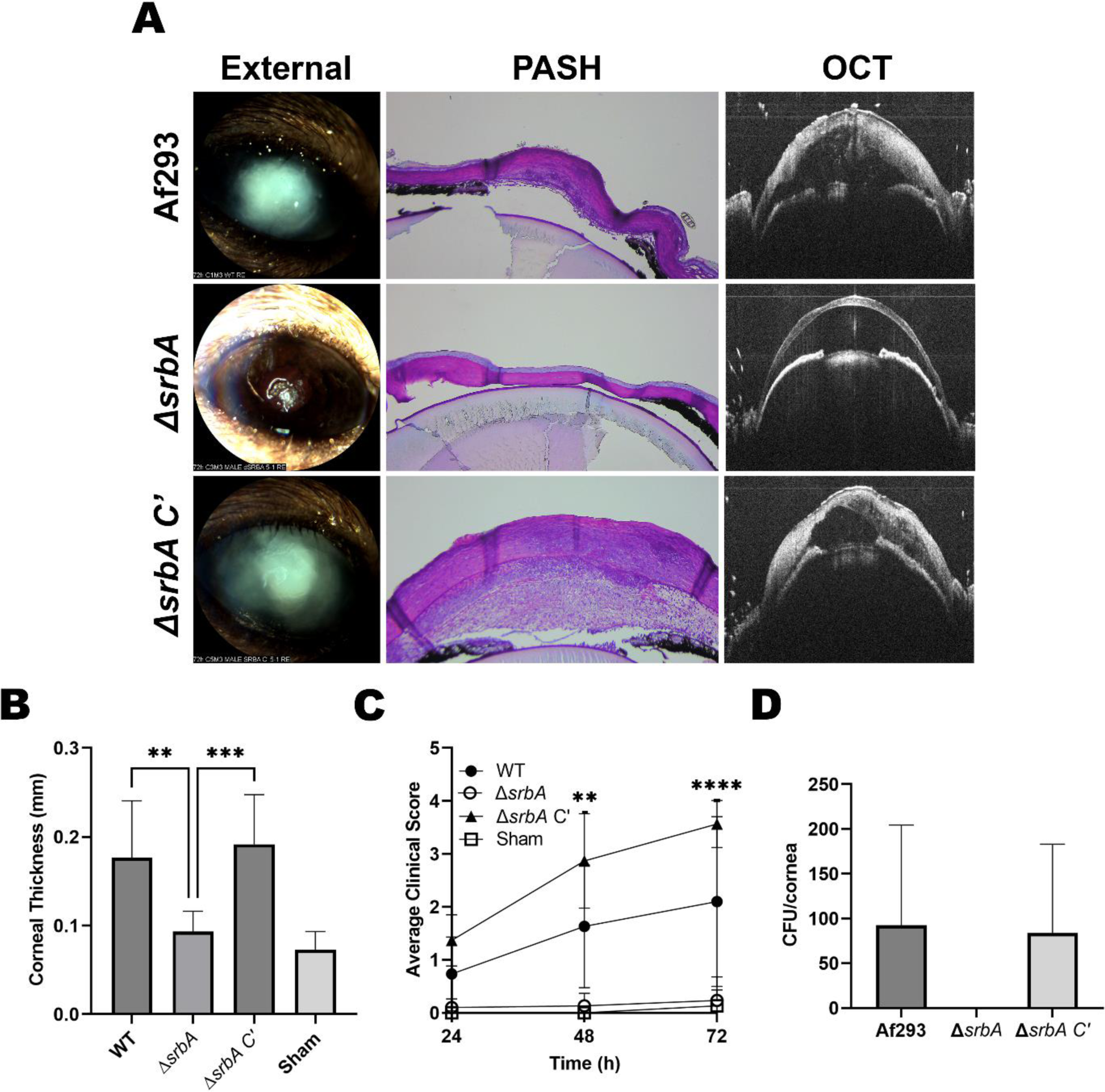
SrbA is essential for virulence in the murine cornea. (A). In corneas infected with Δ*srbA* there was no fungal burden and significantly less inflammation observed by PASH staining at 72 hours p.i. compared to corneas infected with the wild-type or complemented strain (Δ*srbA C’*). (A and B) The Δ*srbA* corneas also maintained their transparency and architecture (measured by OCT) as compared to those infected with the wild-type or the complementor which became edematous and swollen at 72 hours p.i. (C) Disease severity (measured by clinical scoring of all Micron IV images taken every 24 hours after infection) was reduced in corneas infected with Δ*srbA* at 48 and 72 h p.i. (D) No CFU’s were measured from Δ*srbA* infected corneas at 72 h p.i. Statistical Analysis by One-way ANOVA: ** P≤0.01, *** P≤0.001, **** P≤0.0001

## Discussion

Despite its proximity to ambient air, the cornea can become hypoxic downstream to pathologies that increase oxygen consumption by corneal cells or decrease oxygen diffusion through the tissue, such as chemical burn or edema^36–40^. Soft contact lens wear is also a well-described risk factor for corneal hypoxia, which can lead to a breakdown of the epithelial tight junctions that comprise a critical barrier to microbial invasion^41^. Contact lens wear is indeed an important risk factor for FK, suggesting that hypoxia precedes infection in such cases^36,42,43^. The reverse relationship on the other hand is less well understood, i.e., can corneal infection itself be a principal driver of tissue hypoxia? In this study, we demonstrated that oxygen depletion, as assessed through pimonidazole staining, is a prominent feature of two non-contact lens models of *Aspergillus* keratitis in mice. The evaluation of mock-infected corneas suggests that the formation of hypoxia is not related to the predisposing ocular trauma that that precedes inoculation (e.g. debridement or needle puncture), but is likely dependent on the host response to fungal antigen. The potential mechanisms, consequence, and therapeutic implications of these findings will be discussed below.

Rao and Suvas, also using Hypoxyprobe staining, demonstrated that mouse corneal epithelial cells become hypoxic four days following inoculation with Herpes Simplex Virus-1 (HSV-1)^44^. The authors further demonstrate that antibody-mediated neutrophil depletion ablates hypoxia formation, indicating that oxidative metabolism by the inflammatory cells is and underlying cause of oxygen depletion in the model. In our study, we similarly noted an enhanced distribution of pimonidazole staining in FK corneas at time points that correspond with immune cell infiltration (beginning at 48 h p.i.). Notably, however, we also observed diffuse staining across the epithelium and endothelial layers at times that precede clinical or histological signs of inflammation (6 h p.i.). It is known that corneal epithelial cells can respond directly to fungal antigen via Dectin-1 and TLR signaling pathways, leading to NF-kB activation and the secretion of pro-inflammatory cytokines and chemokines^45–50^. It has furthermore been described in other cell types (e.g. adipocytes, retinal pigmented epithelial cells and macrophages) that mitochondrial activity and ROS production are increased shortly after exogenous treatment with such cytokines^50–52^. Thus, we propose that a direct response of epithelial cells to the fungus in our topical inoculation model leads to autologous and paracrine signaling that promotes rapid oxygen depletion across the tissue. When the direct fungal-epithelial interaction is largely bypassed, such as in our intrastromal injection model, we predict that stromal phagocytes (e.g. macrophages, dendritic cells and keratocytes) initiate the cytokine release that then diffuses to both the epithelial and endothelial layers^32^. Future directions will be focused on (1) assessing corneal cell metabolism in response to fungal antigen and pro-inflammatory cytokines and (2) identifying upstream signaling events that promote the metabolic shift in response to fungal antigen.

Numerous studies have shown that the function of all corneal cell types can be altered in low oxygen environments^41,53^. We therefore propose that corneal hypoxia is not only a consequence of FK pathology, but also drives key features of the disease. In our topical model, for example, we observed that the epithelial ulcer fails to reform in infected corneas, whereas it does so by 24 h in sham-inoculated controls (**Figure 1** and data not shown). While direct fungal-mediated damage to the epithelial cells cannot be ruled out as a primary cause of this phenomenon, previous studies have shown that hypoxia decreases the proliferation rate of the basal epithelial cell layer and causes delay in epithelial wound healing^54,55^. Several mechanisms may account for this, including the hypoxia-mediated activation of the polo-like-kinase 3 (Plk3) leading to cell-cycle arrest, as well as a disruption of Ca^2+^ signaling from corneal nerves to epithelial cells ^38,39^. As epithelial ulcers also develop in later stages of our intrastromal model as well as in FK patients, we suggest this may be initiated or exacerbated by the influence of hypoxia on epithelial cell biology^56,57^. Low oxygen may also impact important inflammatory signaling events in the epithelium that influence fungal clearance. Leal *et al,* for example, demonstrated that (1) TLR4 deficiency results in increased fungal (*A. fumigatus*) burden relative to infected wild-type controls, and (2) TLR4^-/-^ mice do not display defects in neutrophil recruitment during infection, suggesting that the TLR4 pathway has a specific role in promoting the fungicidal activity of these cells once they are recruited to the cornea^50^. Importantly, Hara and colleagues showed that human corneal epithelial cells cultured under hypoxic conditions display reduced TLR4 expression^58^. Therefore, the development of hypoxia during FK, which we propose is principally driven by the pro-inflammatory response, may ultimately feedback and inhibit the antifungal activity of inflammatory cells.

The first signs of clinical disease in our FK mice include the development of corneal opacification and increased tissue thickness at around 48 h p.i. Both of these pathologies are consistent with corneal edema, which can result as a breakdown in the barrier and water pumping function of the endothelial layer. In addition to aging and various forms of corneal dystrophy (e.g. Fuchs’ dystrophy), contact lens-associated hypoxia is also known to damage endothelial cell function due to a reduction in ATP production and pump efficiency^59–61^. We therefore reason that the early development of tissue hypoxia may initiate endothelial dysfunction during FK. As corneal edema slowly progresses, so too will the degree of hypoxia, thus forming a feedforward loop of progressive endothelial decline and worsening disease development. At the later time points, it is likely that accumulating fungal metabolites as well as leukocyte-generated ROS/cytokines further drive endothelial cell death, ultimately leading to a critical loss in endothelial cell integrity and a massive influx of fluid from the anterior chamber.

Although corneal edema is associated with a transient increase in corneal thickness measurement, the end-stage consequence of FK in the clinic is often a critical degradation (thinning) of the stromal matrix that results in perforation and a need for corneal transplantation^62,63^. This breakdown in the collagen fibers is likely attributable to secreted proteases from both the fungus as well as matrix metalloproteases (MMPs) secreted by the corneal fibroblasts and infiltrating leukocytes^6,64–67^. Regarding corneal fibroblasts, it has been demonstrated that hypoxic culture increases MMP-1 and MMP-2 production, while the production of various collagen species decreases. Indeed, stromal thinning is a consequence of contact-lens driven hypoxia^68–70^. Similarly, hypoxia has been shown to increase MMP-9 from neutrophils, along with other collagen damaging molecules including elastase, and myeloperoxidase^71^. Taken together, the development of hypoxia during FK likely promotes the progression of numerous disease pathologies spanning all cellular layers of the cornea, and the intervention of key cellular regulators of these phenomena (e.g. HIF-1) may play a protective role.

Our studies demonstrate that the development of hypoxia across the epithelium is not only rapid, but strong enough to render the *A. fumigatus* SrbA essential for the establishment of invasive growth into cornea. This differs somewhat from previous studies in the lung, whereby the Δ*srbA* mutant is able to initiate growth (germinate) in the airway, but fails to maintain its growth and virulence potential after the onset of neutrophil-driven tissue inflammation at around 2 days post-inoculation. Thus, while *A. fumigatus* SrbA is a critical determinant of fungal virulence and disease outcome in both IPA and FK, the stage of infection at which hypoxia develops and renders SrbA essential differs between the lung and corneal environments. One key difference may be the high degree of vascularization of the airway, which oxygenates the tissue and buffers the development of hypoxia that would otherwise be driven by the pulmonary epithelial response to fungal antigen.

Azoles are a major class of antifungals for both invasive and ocular *Aspergillus* infection^72–77^. These drugs target a key enzyme in the sterol biosynthetic pathway and, in this way, deplete membrane sterol content in the same way hypoxia does. As such, SrbA is activated upon azole treatment, and the *ΔsrbA* mutant is hypersensitive to several drugs in this class, including fluconazole, to which wild-type *A. fumigatus* is resistant (**Figure S2C**)^20^. Thus, pharmacological inhibitors, if they could be developed, of the *A. fumigatus* SrbA pathway would not only be inherently antifungal in hypoxic tissues, but they could potentiate the efficacy of an already available class of antifungals. Ongoing studies in our group are aimed at identifying key regulators of hypoxia adaptation in *Fusarium* species, which as we have shown here, is likely an important aspect of virulence in this pathogen as well.

In summary, and to our knowledge, this study is the first to demonstrate an important role for corneal hypoxia in the pathobiology of fungal keratitis. While our data directly demonstrate an influence of hypoxia on the fungus, it is likely that oxygen depletion will alter the function of effectively all host cells and influence disease pathogenesis. A better understanding of how hypoxia influences the host-fungus interaction, as well as how other FK-relevant pathogens adapt to hypoxia, will undoubtedly give insights into novel therapeutic interventions that improve patient outcomes in FK patients.

## Methods

### Strains and culture conditions

Strains and plasmids used in this study are listed in **Table S1**. All *A. fumigatus* strains were derived from Af293 and maintained on Glucose Minimal Medium (GMM), (adapted from Cove 1966), containing 1% (w/v) Glucose and 10 µM ammonium tartrate as a nitrogen source. Cultures grown in hypoxia were grown in 1% O_2_ and 5% CO_2_. All incubations were performed at 35°C. *F. solani* was maintained on Potato Dextrose Agar and incubated at 30°C.

### Generating mutants

For our mutants, *Aspergillus fumigatus* strain Af293 expressing mCherry from an ectopic integration (Af293 *PgpdA-mCherry-hph*) was used as the wild type (WT) organism. Targeted gene deletions were performed as described in Al Abdallah *et al* 2019 and Szewczyk et al 2006^78,79^. Protoplasts were generated from germlings using the following enzymes for 3.5 hours: 5 mg/ml of Lysing Enzymes from *Trichoderma harzanium*, 5 mg/ml of Driselase, and 100 µg of chitinase in OSM (1.2 M MgSO_4_, 10 mM Sodium Phosphate Buffer pH 5.8) at 30 °C. Protoplasts were purified using a density-based trapping buffer (0.6M Sorbitol, 0.1 M Tris-HCl) layered over the Lysing Enzyme Buffer and spun at 5000 x g for 15 minutes at 4 °C. The protoplasts were pipetted from the interphase, washed via centrifugation, 5 minutes at 5000 x g, in STC (1.2 M sorbitol, 10mM CaCl_2_, 10 mM Tris-HCL pH 7.5), enumerated via hemocytometer, and diluted to 5 x 10^5^ protoplasts/ml.

Crispr RNAs (cRNAs) and primers used in this study are listed in **Table S2**. crRNAs were designed to flank the coding region of the *srbA* gene (Afu2g01260) and conjugated to tracrRNAs (IDT) by combining equimolar amounts with Nuclease Free Duplex Buffer (IDT). 3.33 μl of each reagent is combined and incubated at 95 °C for 5 minutes before cooling them to room temperature. The crRNAs were then conjugated to 1μg of Alt R Cas9 (IDT) in 10 μl of Cas9 Working Buffer (20 mM HEPES, 150 mM KCl pH of 7.5) at 37 °C for 30 minutes. 200 μl of the protoplasts were transferred to a fresh Eppendorf tube and mixed with 10 μg of the phleomycin repair template (amplified from p402R), flanked with 35bp regions of microhomology directly adjacent to the Cas9 mediated double stranded breaks. The Cas9 conjugated gRNAs were combined and introduced to the protoplasts. 50 μl of PEG solution (60% PEG M.W. 3350 [w/v], 50 mM CaCl_2_, 50 mM Tris-HCL pH 7.5) was added to the protoplast/DNA mixture and mixed by flicking the tube. The mixture was then placed on ice for 50 minutes. An additional 500 μl of the PEG solution was added and incubated at room temperature for 30 minutes. The PEG solution was removed by centrifuging the protoplasts at >13,000 rpm for 3 minutes and pipetting off the supernatant. The protoplasts were centrifuged at >13,000 for an additional 1 minute to remove the remaining PEG solution before they were resuspended in 1 ml of STC. 100 μl of this suspension was plated onto 20ml of GMM + 1.2M Sorbitol and incubated at room temperature overnight. A 10 ml overlay of Top Agar (GMM with 5g/L of Agar) and 375 μg/ml phleomycin was poured onto the protoplasts. These plates were placed at 35 °C and monitored for the formation of microcolonies for 48 hours. Colonies that grew under selection were screened via PCR for a replacement of the *srbA* coding region with the phleomycin resistance marker *bleR*.

*srbA* was complemented into the *atf4* locus using a single crRNA mediated cut site and a marker-less transformation strategy^35^. Protoplasts were recovered in hypoxia (1% O_2_, and 5% CO_2_) after an overnight incubation at room temperature and ambient oxygen conditions. Colonies that were able to grow in hypoxia were then screened via PCR for the correct insertion of the *srbA* locus into the *atf4* locus.

### Topical Murine Keratitis model

These experiments were preformed according the Association for Research in Vision and Ophthalmology guidelines for the use of animals in vision research. Groups of C57BL/6J mice were obtained from Jackson Laboratories. On the day preceding fungal infection, the animals were immunosuppressed via intraperitoneal injection with 100 mg/kg of methylprednisolone. On the day of infection, conidia were swollen in YPD (10% Yeast Extract, 20% Peptone, 20% Glucose) for 4.5 hours at 35 °C to ensure that the conidia are in isotropic growth and were metabolically active. The inoculum was normalized to an OD 360nm. Mice were anesthetized with Ketamine (100mg/kg) and Xylazine (6.6mg/kg). Once the mice were under surgical anesthesia, the center of the right cornea was Algerbrushed to remove the epithelium, and 5μl of the inoculum was placed onto the ulcerated cornea and allowed to rest on the ocular surface for 20 minutes before being wicked off with a Kim wipe. Buprenorphine SR (1mg/kg) was administered subcutaneously intraoperatively for analgesia.

### Intrastromal Murine Keratitis model

These experiments were preformed according the Association for Research in Vision and Ophthalmology guidelines for the use of animals in vision research. Infections were performed as described in Leal et al 2014^28^. Briefly, mice were immunosuppressed with 100 mg/kg of methylprednisolone on the day preceding infection. On the day of infection, the mice were anesthetized with Ketamine and Xylazine intraperitoneally. A small pocket was introduced into the corneal epithelium with a 20.5-gauge needle before a glass-pulled capillary needle was inserted into the opening. Using a programmable pneumatic microinjection system (Microdata Instruments, Plainfield, NJ, USA) 2 μl of 2.5×10^7^ conidia/ml were injected into the corneal stroma. Buprenorphine SR was administered subcutaneously intraoperatively for analgesia.

### CFU analysis

Corneas were resected and placed in 2 mg/ml of Collagenase I (Sigma SCR103) in PBS within a 1.5 ml Eppendorf Tube. The tubes were oscillated at 50 Hz in two 30 second pulses in a Qiagen TissueLyzer before they were incubated at 37 °C for 15 minutes. The homogenate was then pipetted vigorously 8 times before being returned to 37 °C for an additional 15 minutes. These steps were repeated once more before the homogenate was diluted 1:1 in PBS and 100 μl was plated onto Inhibitory Mold Agar (IMA). Plates were incubated overnight at 35 °C before colonies were enumerated.

### Micron IV Imaging and Analysis

Mice were anesthetized with isoflurane and slit lamp images were taken using a Micron IV Biomicroscope (Phoenix Technology Group CA, USA). Bright field images were captured at a 66ms exposure.

### OCT Imaging and Analysis

Corneas were imaged using the Bioptigen spectral-domain-optical coherence tomography system (Leica Microsystems, Buffalo Grove, IL) to measure corneal edema and inflammation. Mice were anesthetized with isoflurane and a 4×4 mm image is scanned with the 12mm telecentric lens. Reference arm calibration was completed by the manufacturer and set to 885.

Corneal scans were digitally overlaid with an 11×11 Spider plot, and the 13 central points were marked at the endothelium and the epithelium. The differences between these points were calculated as the corneal thickness in millimeters. These 13 measurements were then averaged and plotted as a single data point.

### Hypoxyprobe Immunohistochemical staining

Mice received 60 mg/kg of Hypoxyprobe, dissolved in PBS, 90 minutes before they were euthanized. Eyes were enucleated and fixed in paraformaldehyde at the Dean McGee Histology Core. To deparaffinize the sections the slides were warmed up in a 40°C oven for 20 minutes. The slides were then placed in a xylene bath for 5 minutes three times. Sections were rehydrated in three separate 95% ethanol baths three times before being washed in washing buffer (0.2% (w/v) Brij 35 in PBS)

Slides were incubated at 90°C for 20 minutes in High pH IHC Antigen Retrieval Solution (Thermo Fisher Scientific Catalog Number 00-4956) and allowed to cool to room temperature. Slides were then washed and blocked with Cas-Block (Life Technologies 00-8120) for 120 minutes. Sections were incubated with the primary antibody, 1:100 FITC-Mab1 (Hypoxyprobe mouse MAb clone 4.3.11.3) overnight at room temperature.

Slides were then washed and incubated for 60 minutes at room temperature with the secondary antibody: 1:100 dilution of DyLight 594 conjugated mouse anti-FITC (Jackson Immunoresearch 200-512-037).

Slides were washed and dried by blotting before being mounted and counterstained using ProLong Gold Antifade Mountant with DAPI (ThermoFisher P36931).

Imaging was performed on a Nikon E800 Epiflourescent Microscope. Single color channel images were captured in DAPI, and Dylight 594. Exposure times were 3 seconds for DAPI and 5 seconds for Dylight 594. Images were processed in Metamorph (Molecular Devices, San Jose, CA) and background fluorescence was determined for each single channel using isotype controls.

### Statistical Analysis

Statistical analysis performed on GraphPad Prism 10.0.2

---

**Supplementary Figure 1.**
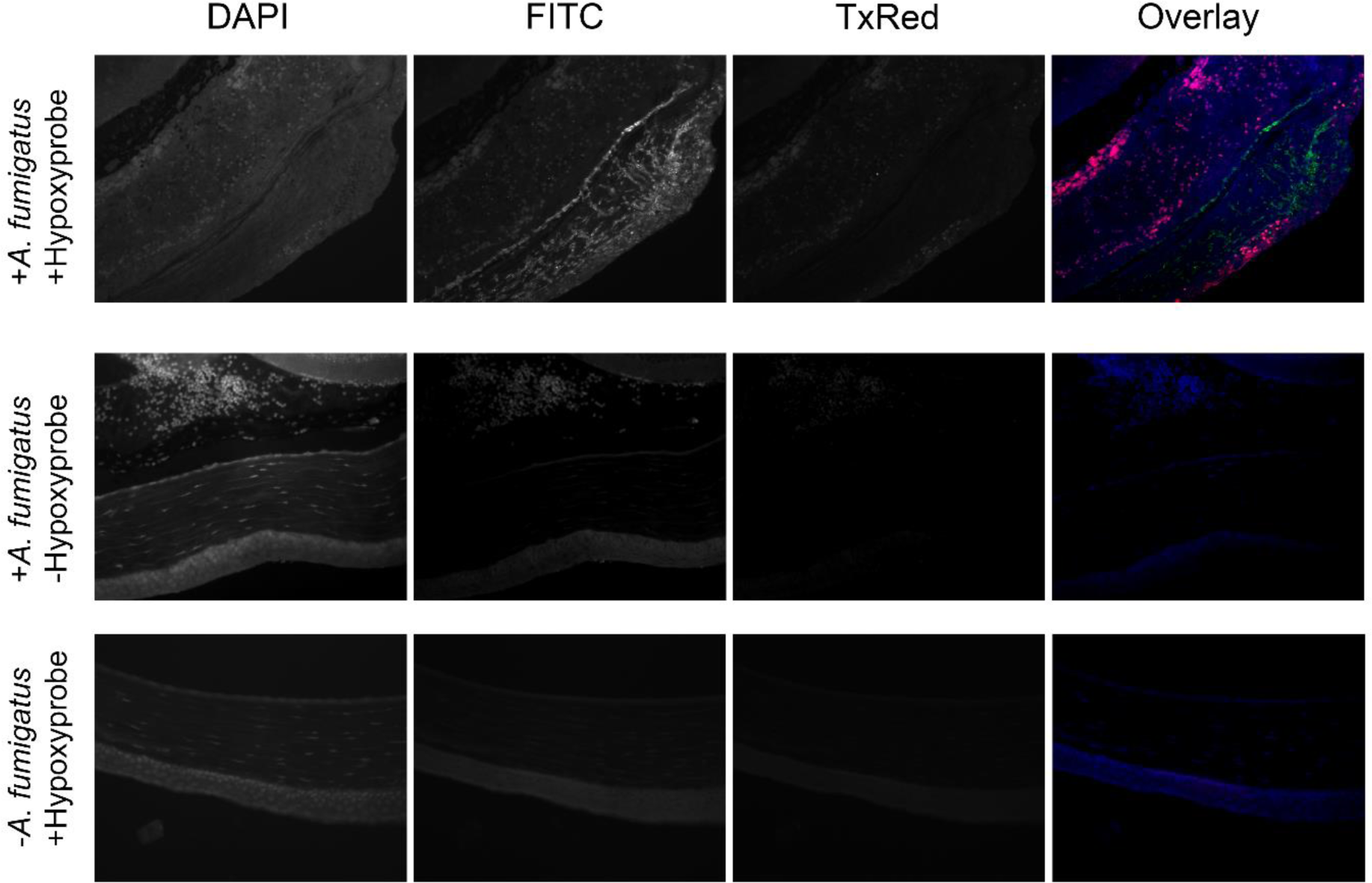
Isotype controls for Hypoxyprobe show specific staining. Sections of infected tissue from mice not injected with Hypoxyprobe show no positive Hypoxyprobe signal. Healthy tissues show no Hypoxyprobe signal above background threshholds. Single color channel pictures were merged to show overlapping localization of corneal epithelium (DAPI, blue), fungal burden (green), and Hypoxyprobe (red).

**Supplementary Figure 2.**
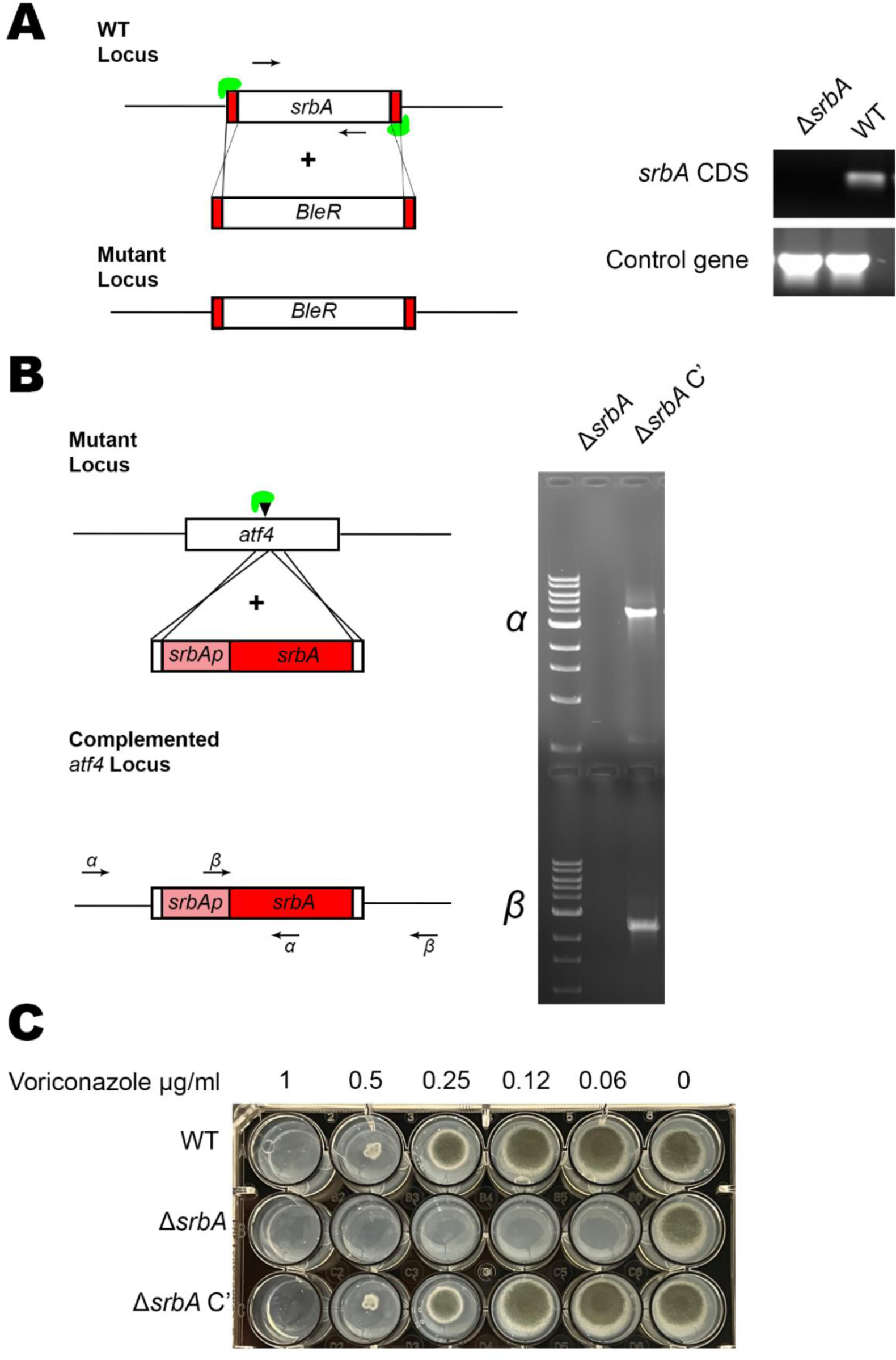
Generation of Δ*srbA* mutant and complemented strain (Δ*srbA C’*). (A) An *in vitro* assembled Cas9 RNP approach was used to generate the Δ*srbA* knockout mutant. Mutants were confirmed by PCR (B) The *srbA* gene and its native promoter were amplified with primers containing microhomology sequences to the *atf4* safe haven locus. The *in vitro* Cas9 approach was used to make a single cut in the *atf4* locus and the *srbA* cassette insertion was confirmed by PCR. (C) The Δ*srbA* mutant, the wild-type (WT), and the complemented strain (Δ*srbA* C’) were grown at varying concentrations of voriconazole in Glucose Minimal Media at 35°C and imaged at 72 h post-inoculation. The Δ*srbA* mutant is hypersensitive to azoles.

**Supplementary Table 1.**
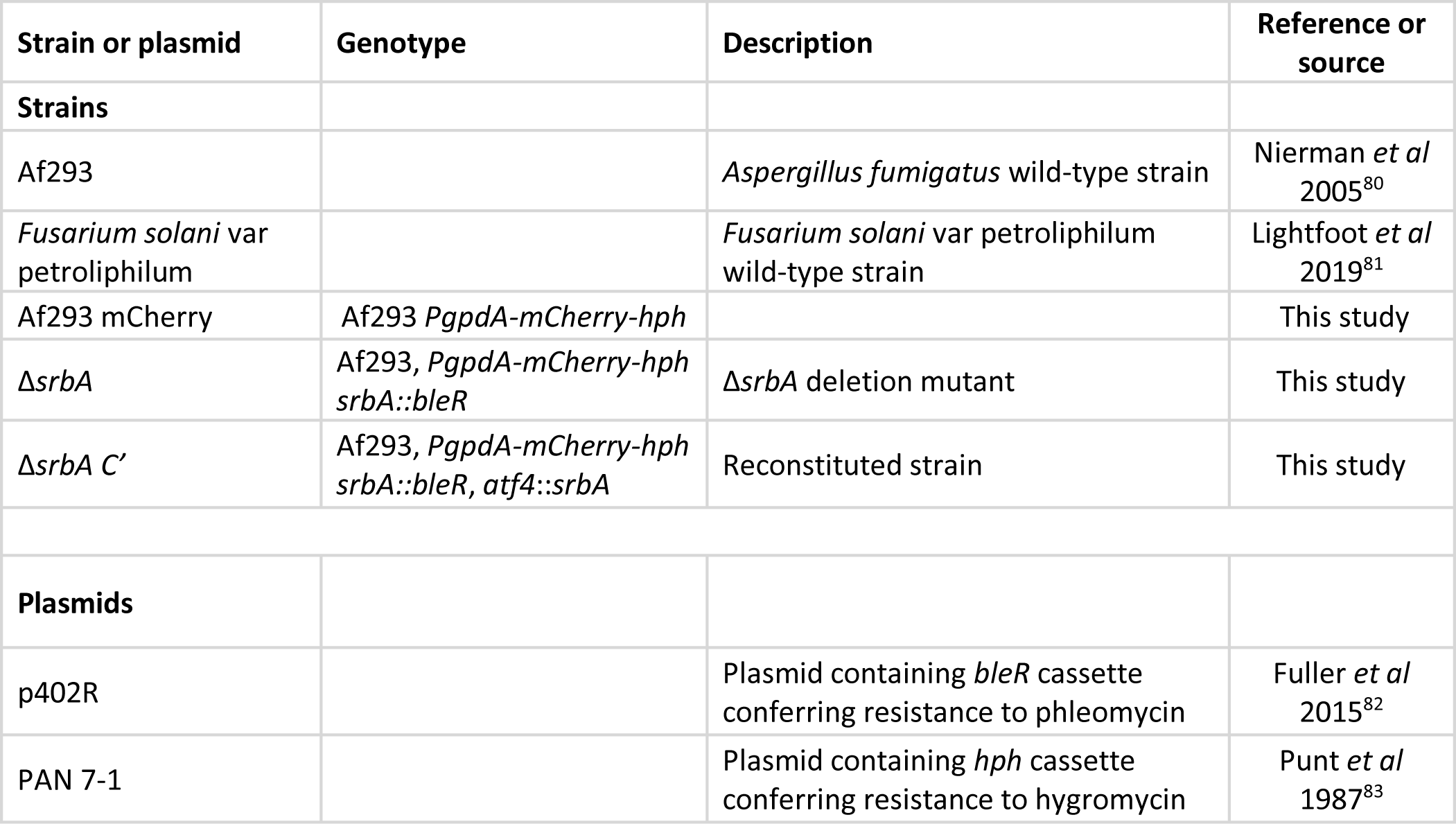
Strains and Plasmids used in this study.

| Strain or plasmid | Genotype | Description | Reference or source |
| --- | --- | --- | --- |
| **Strains** |  |  |  |
| Af293 |  | *Aspergillus fumigatus* wild-type strain | Nierman *et al* 2005<sup>80</sup> |
| *Fusarium solani* var petroliphilum |  | *Fusarium solani* var petroliphilum wild-type strain | Lightfoot *et al* 2019<sup>81</sup> |
| Af293 mCherry | Af293 *PgpdA-mCherry-hph* |  | This study |
| Δ*srbA* | Af293, *PgpdA-mCherry-hph*<br>*srbA::bleR* | Δ*srbA* deletion mutant | This study |
| Δ*srbA C'* | Af293, *PgpdA-mCherry-hph*<br>*srbA::bleR, atf4::srbA* | Reconstituted strain | This study |
| **Plasmids** |  |  |  |
| p402R |  | Plasmid containing *bleR* cassette conferring resistance to phleomycin | Fuller *et al* 2015<sup>82</sup> |
| PAN 7-1 |  | Plasmid containing *hph* cassette conferring resistance to hygromycin | Punt *et al* 1987<sup>83</sup> |

**Supplementary Table 2.**
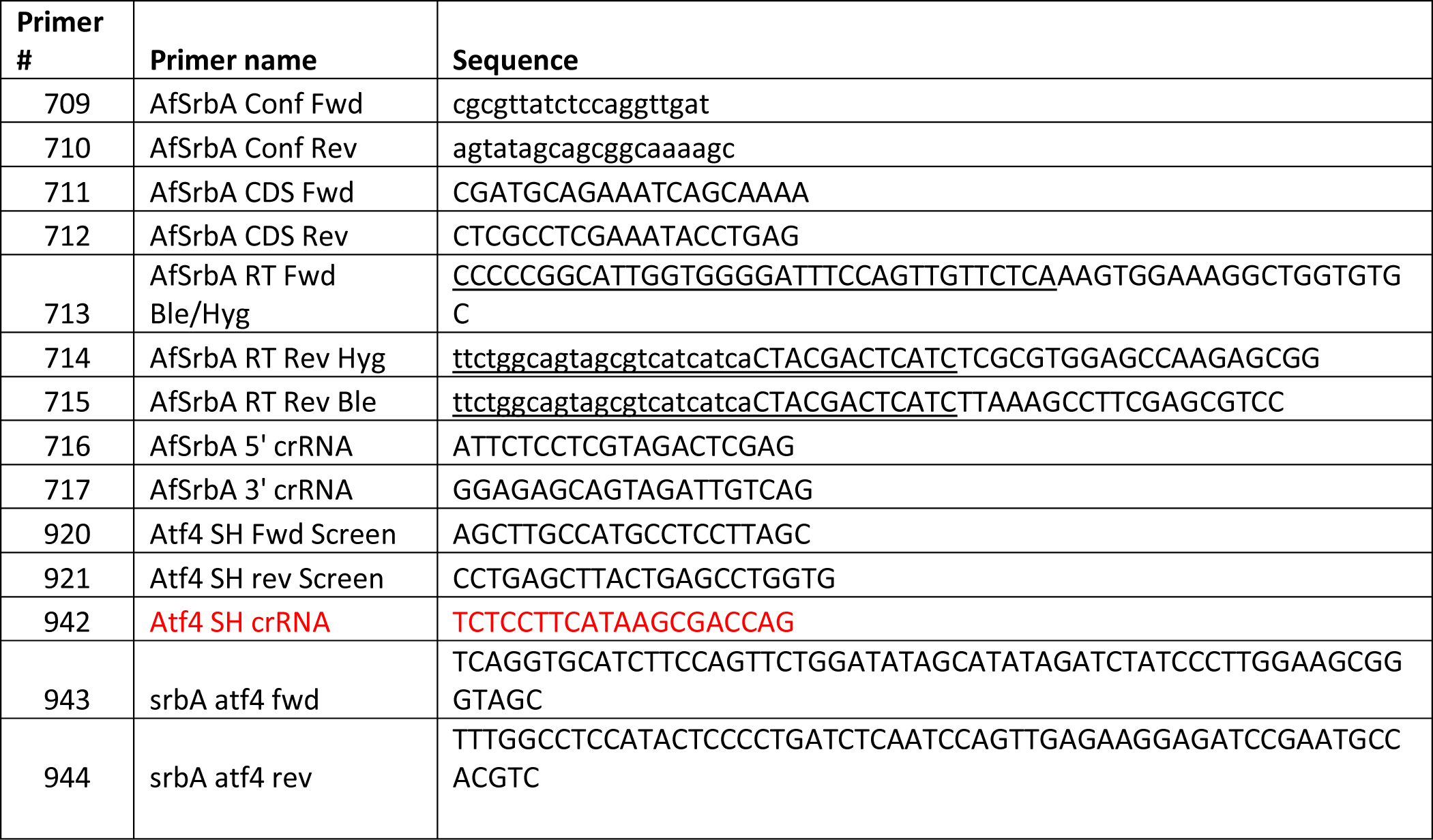
Nucleotide Sequences for primers and crRNA’s used for deletion and complementation strain construction.

